# Stable High-Concentration Monoclonal Antibody Formulations Enabled by an Amphiphilic Copolymer Excipient

**DOI:** 10.1101/2022.05.25.493499

**Authors:** John H. Klich, Catherine M. Kasse, Joseph L. Mann, Yaoqi Huang, Andrea I. d’Aquino, Abigail K. Grosskopf, Julie Baillet, Gerald G. Fuller, Eric A. Appel

**Affiliations:** Department of Bioengineering, Stanford University, Stanford, CA 94305, USA; Department of Materials Science & Engineering, Stanford University, Stanford, CA 94305, USA; Department of Chemical Engineering, Stanford University, Stanford, CA 94305, USA; Department of Pediatrics – Endocrinology, Stanford University School of Medicine, Stanford, CA 94305, USA; ChEM-H Institute, Stanford University, Stanford, CA 94305, USA; Woods Institute for the Environment, Stanford University, Stanford, CA 94305, USA

**Author notes:** These authors contributed equally to this work (John H. Klich, Catherine M. Kasse).

**Keywords:** Antibodies, Formulation, Biopharmaceutical, Excipient, Drug Delivery

## Abstract

Monoclonal antibodies are a staple in modern pharmacotherapy. Unfortunately, these biopharmaceuticals are limited by their tendency to aggregate in formulation, resulting in poor stability and often requiring low concentration drug formulations. Moreover, existing excipients designed to stabilize these formulations are often limited by their toxicity and tendency to form particles such as micelles. Here, we demonstrate the ability of a simple “drop-in”, amphiphilic copolymer excipient to enhance the stability of high concentration formulations of clinically-relevant monoclonal antibodies without altering their pharmacokinetics or injectability. Through interfacial rheology and surface tension measurements, we demonstrate that the copolymer excipient competitively adsorbs to formulation interfaces. Further, through determination of monomeric composition and retained bioactivity through stressed aging, we show that this excipient confers a significant stability benefit to high concentration antibody formulations. Finally, we demonstrate that the excipient behaves as an inactive ingredient, having no significant impact on the pharmacokinetic profile of a clinically relevant antibody in mice. This amphiphilic copolymer excipient demonstrates promise as a simple formulation additive to create stable, high concentration antibody formulations, thereby enabling improved treatment options such as a route-of-administration switch from low concentration intravenous (IV) to high concentration subcutaneous (SC) delivery while reducing dependence on the cold chain.

## 1. Introduction

Biopharmaceuticals are rapidly replacing small organic molecules as the leading active pharmaceutical ingredients (APIs) on the market today.^[1]^ Monoclonal antibodies (mAbs) are one broadly important class of biopharmaceuticals that have been thoroughly investigated for their superior target selectivity and reduced toxicity when compared to small organic molecules. These advantages notwithstanding, many antibody-based therapeutics have been limited in their translational success due to formulation instabilities, most generally aggregation, arising from their complex tertiary structure, large molecular weight, and inherently amphiphilic nature.^[2, 3]^ While the mechanism of aggregation varies slightly from antibody to antibody, many contain hydrophobic patches or partially denature to reveal hydrophobic segments of the molecule that drive adsorption of the proteins onto interfaces (e.g., air-water, glass-water, or rubber-water interfaces).^[3-5]^ The high local concentrations of the molecules at these interfaces, coupled with interface-mediated partial unfolding, trigger initial nucleation of aggregation events that lead to further aggregation.^[3, 5]^ This tendency to aggregate at interfaces increases with increasing formulation concentration, negatively impacting overall formulation stability. These inherent concentration limitations often require that large-volume, low-concentration transfusions of mAb (monoclonal antibody) therapies are delivered intravenously (IV) (Figure 1A). Unfortunately, IV administration places a larger burden on patients compared to other routes of administration, requiring lengthy transfusion procedures and access to clinical infrastructure, which at best is inconvenient and at worst could preclude patients from receiving effective treatment. High concentration formulations of mAbs would be necessary to enable dose-matched subcutaneous (SC) injections that are significantly less burdensome on patients and which can also be administered in low resource settings (Figure 1A).^[6]^

**Figure 1.**
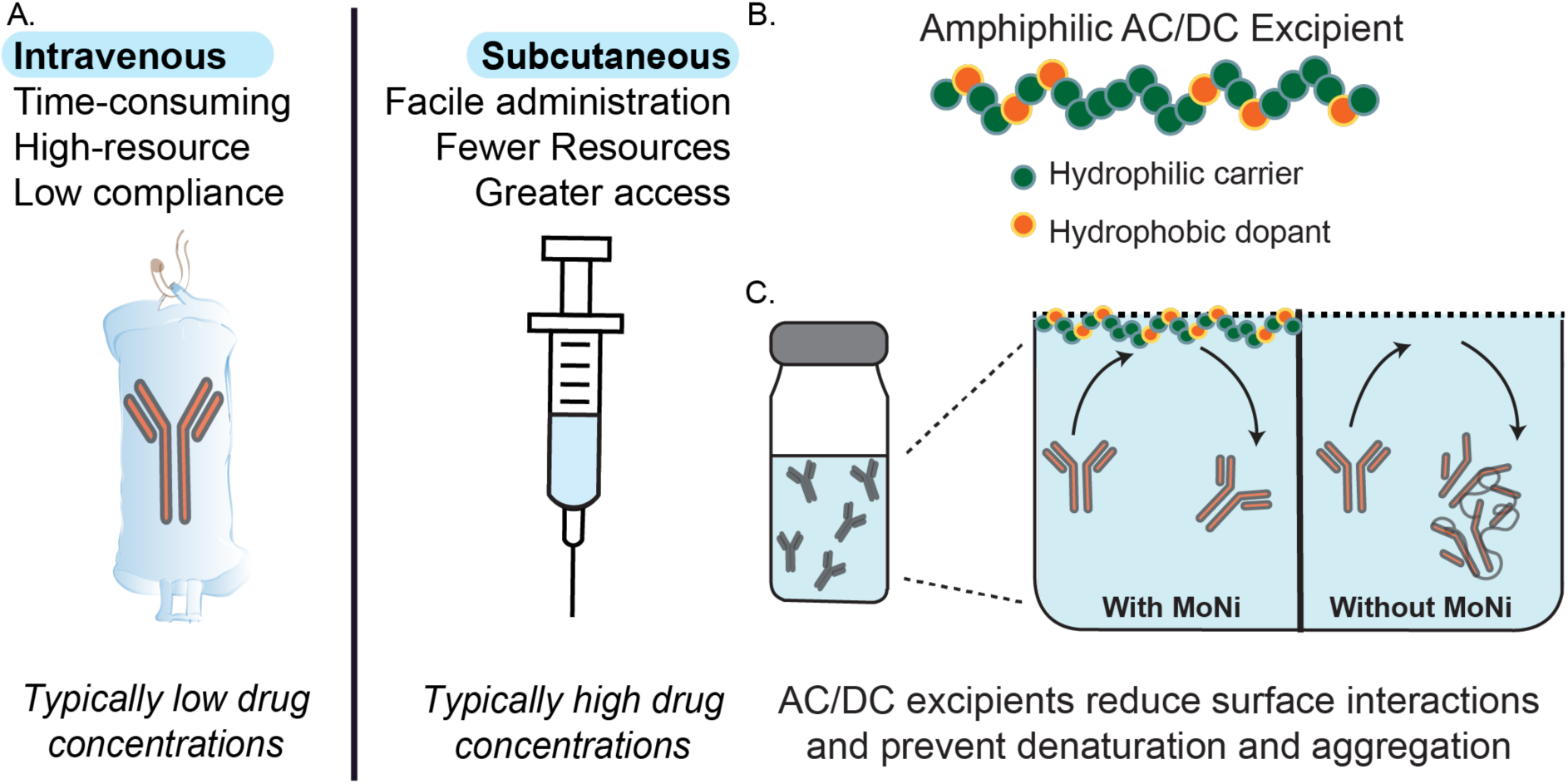
Overview of stabilization of high-concentration antibody formulations using novel copolymer excipients. (A) Schematic overview of current therapeutic routes of administration for biopharmaceuticals, whereby intravenous administration is common because of limitations on stability of biopharmaceutics at the high concentrations required for subcutaneous administration, resulting in high patient burden. (B) Graphical representation of amphiphilic acrylamide carrier/dopant copolymer (AC/DC) excipients, such as lead candidate poly(acryloylmorpholine-co-*N*-isoproylacrylamide) (MoNi), that can be used to stabilize biopharmaceuticals in formulation. (C) Proposed mechanism of stabilization of monoclonal antibodies in formulation using AC/DC copolymers as simple “drop-in” excipient. Since AC/DC copolymers are amphiphilic, they preferentially absorb to interfaces, precluding antibody interfacial adsorption in formulation and thereby preventing interface-related destabilization and aggregation events.

Formulation limitations are particularly detrimental for clinical indications that require maintenance of high mAb serum concentrations for therapeutic benefit. For example, broadly neutralizing HIV antibodies (bnAbs) are a class of antibody therapeutics specifically targeting highly conserved regions on the HIV envelope trimer.^[7]^ HIV bnAbs are currently being explored for their use in prophylactic infection prevention and disease management through passive immunization.^[8]^ While the use of bnAbs for HIV prevention and treatment has been demonstrated in mice and non-human primates,^[9]^ bnAb formulations must typically be administered by IV infusion to reach the high serum concentrations required for protection against breakthrough infection and to prevent onset of drug resistance.^[8]^ According to UNAIDS, there were 1.7 million new annual infections and approximately 38 million people living with HIV in 2021; it follows that the development of new therapeutics for the treatment and prevention of HIV that can be deployed even in resource limited settings remains a high priority. SC mAb therapy is a more scalable pathway towards providing prophylactic protection to at-risk populations and treatment for those currently living with HIV, but as noted, this strategy necessitates the development of stable, high concentration mAb formulations to simplify treatment and improve global access.^[6, 10][11]^

The addition of excipients to biopharmaceutical formulations is a common strategy to overcome challenges associated with high concentration formulation of mAbs. Biopharmaceutical formulations commonly contain polyoxyethylene-based surfactants such as polysorbates and poloxamers.^[12, 13]^ Despite their widespread use, these systems are still rather limited by their critical micelle concentrations (the concentration above which the excipient will form micelles, further destabilizing the formulation through introducing large hydrophobic surfaces), possible toxicity through oxidative degradation, and undesirable interactions between the excipient and the cargo in the bulk.^[13]^ A more translatable approach towards stabilizing biopharmaceuticals would use an excipient that could be easily and stably incorporated into highly concentrated commercial formulations without interacting with the therapeutic in the bulk, altering its pharmacokinetics, or increasing toxicity.

A promising strategy to stabilize protein formulations is using amphiphilic acrylamide carrier/dopant copolymer (AC/DC) excipients, which preferentially occupy interfaces to prevent interface-mediated protein aggregation. In previous work, we have used this class of excipients to stabilize low molecular weight (<5.8 kDa) biopharmaceuticals such as insulin analogues and pramlintide (an amylin analogue).^[14, 15]^ Further, we previously demonstrated that these AC/DC excipients are biocompatible and exhibit low toxicity.^[14, 15]^ However, these excipients have not yet been demonstrated to stabilize mAbs at high concentrations in formulation, where the active pharmaceutical ingredient is both much larger (150 kDa compared to 5.8 kDa) and formulated at much higher concentrations (>100 mg/ml compared to 3.5 mg/ml).

In this study, we utilize a lead-candidate AC/DC excipient, poly(acryloylmorpholine-co-*N*-isopropylacrylamide) (MoNi), to enable high-concentration formulation of PGT121, a promising anti-HIV bnAb, that is suitable for SC injection. Mechanistically, we investigate the ability of MoNi to preferentially occupy the interfaces in mAb formulations using both interfacial rheology and surface tension measurements. Next, we probe the stability of PGT121 formulations with and without added MoNi using accelerated aging assays at different API concentrations. We then conduct pharmacokinetic studies in mice, comparing SC administration of a low volume of a MoNi-stabilized high-concentration PGT121 formulation to intraperitoneal (IP) administration of a larger volume of a standard low-concentration PGT121 formulation. Overall, we demonstrate that MoNi significantly improves the stability of mAb formulations without altering pharmacokinetics or bioavailability, illustrating its potential as a “drop-in” excipient.

## 2. Results and Discussion

### 2.1. AC/DC excipient selectively adsorbs to interfaces in mAb formulations

From a previous screen of hundreds of amphiphilic AC/DC copolymers for their ability to stabilize insulin, we identified several top-performing excipient candidates.^[14, 15]^ In particular, a copolymer of acryloylmorpholine and *N*-isopropylacrylamide, referred to as “MoNi”, exhibited exceptional ability to stabilize proteins in formulation by preferentially occupying interfaces.^[15]^ We hypothesized that MoNi could similarly impart improved stability to mAb formulations by reducing interface-mediated changes in protein conformation and aggregation-nucleating events, thereby improving formulation stability (Figure 1B,C).

To synthesize MoNi, we employed reversible addition fragmentation transfer (RAFT) controlled radical polymerization on account of its simple implementation and excellent control over the synthesis of copolymers. Although RAFT polymerization has many synthetic advantages, it generates polymers with a reactive trithiocarbonate chain transfer agent (CTA) attached at the Z terminus of the polymer following polymerization, denoted here as MoNi-CTA. We removed the CTA moiety from the MoNi-CTA copolymer to produce the MoNi excipient before utilization in subsequent assays to ensure both chemical stability and biological inertness (Figure 2A-D). Absence of the absorbance peak in the SEC trace confirms removal of the UV-active Z group (UV_max_/RI_max_ ∼ 7 before cleavage and <0.1 after cleavage; Figure 2C). In previous studies, we have demonstrated that MoNi is highly biocompatible and exhibits exceptionally low cytotoxicity compared to typical surfactant molecules.^[15]^ By comparing reported values for the concentration whereby 50% of cells are killed *in vitro*, a parameter called the LC_50_, MoNi is over 100-fold less cytotoxic than common commercial surfactant excipients such as Polysorbate 80 (PS80) and Pluronic L61 (Figure 2E). The MoNi copolymer excipient is synthesized at molecular weights well below the glomerular filtration threshold to ensure that they are rapidly excreted following administration in the body without bioaccumulation.^[15]^ Further, we attempted to determine MoNi’s critical micelle concentration (CMC) by performing dynamic light scattering (DLS) analysis on a dilution series of MoNi from 100 to 1 mg/mL (Figure 2F). In these assays, the MoNi excipient did not form micelles or aggregates within the concentration range assessed. This behavior is unique compared to the common commercial surfactant excipients PS80 and Pluronic L61, which exhibit CMC values well below 1 mg/mL (Figure 2G). Importantly, most surfactants are toxic in their micellular state, suggesting that the significantly lower cytotoxicity of MoNi may arise from the fact that it does not exhibit self-assembly into micelles at formulation-relevant concentrations.^[17]^

**Figure 2.**
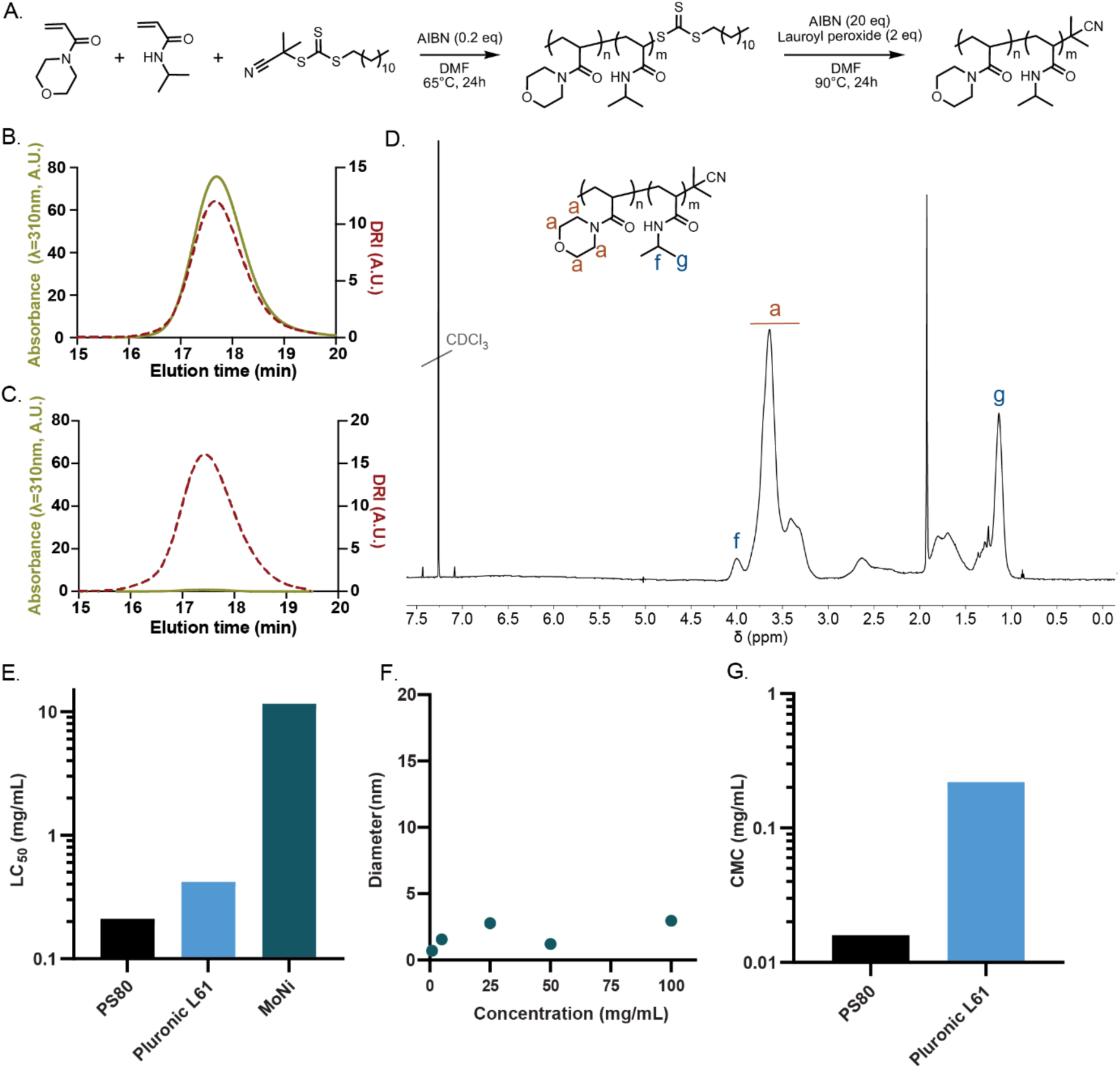
Synthesis and characterization of MoNi copolymer excipient. (A) Synthetic scheme for the synthesis of poly(acryloylmorpholine-co-*N*-isopropylacrylamide) (MoNi). Size exclusion chromatography (SEC) characterization of the (B) MoNi-CTA (chain transfer agent) intermediate and (C) MoNi demonstrates that the CTA is completely removed in the second step of the synthesis, yielding a chemically stable copolymer. (D) ^1^H-NMR characterization of MoNi. (E) Comparison of cytotoxicity of MoNi^[15]^, polysorbate 80^[16]^, and pluronic L61^[17]^. (F) DLS characterization of MoNi in solution at various concentrations. (G) Comparison of critical micelle concentration (CMC) values for polysorbate 80^[18]^, and pluronic L61^[19]^. MoNi did not exhibit micelle formation within the assessed concentration range (100 to 1 mg/mL).

We then sought to evaluate our hypothesis that MoNi preferentially adsorbs to the air-water interface, thereby precluding mAb adsorption to the interface and improving formulation stability by reducing interface-mediated aggregation events. We first conducted time-resolved surface tension experiments with formulations of the anti-HIV bnAb PGT121, a promising candidate for passive immunization against HIV targeting the well conserved V1/V2 glycan on the gp120 surface glycoprotein that is currently being evaluated clinically.^[20, 21]^ We observed that formulations of PGT121 (20 mM acetate buffer, pH∼5.2) with MoNi (0.01 wt%) exhibited lower surface tension values compared to PGT121 alone (53 mN/m and 62 mN/m, respectively; Figure 3A). The lower surface tension observed for the PGT121 formulation comprising MoNi suggests that there are more species packed at the interface in this formulation than in the formulation with only PGT121, indicating that MoNi is preferentially adsorbing to the interface at higher levels. Notably, the surface tension of both a formulation of MoNi only in buffer and a formulation of PGT121 and MoNi were found to be identical (Figure 3A), indicating that the air-water interface is indeed dominated by MoNi at these concentrations. These results suggest that MoNi preferentially adsorbs to the air-water interface, precluding mAb adsorption.

**Figure 3:**
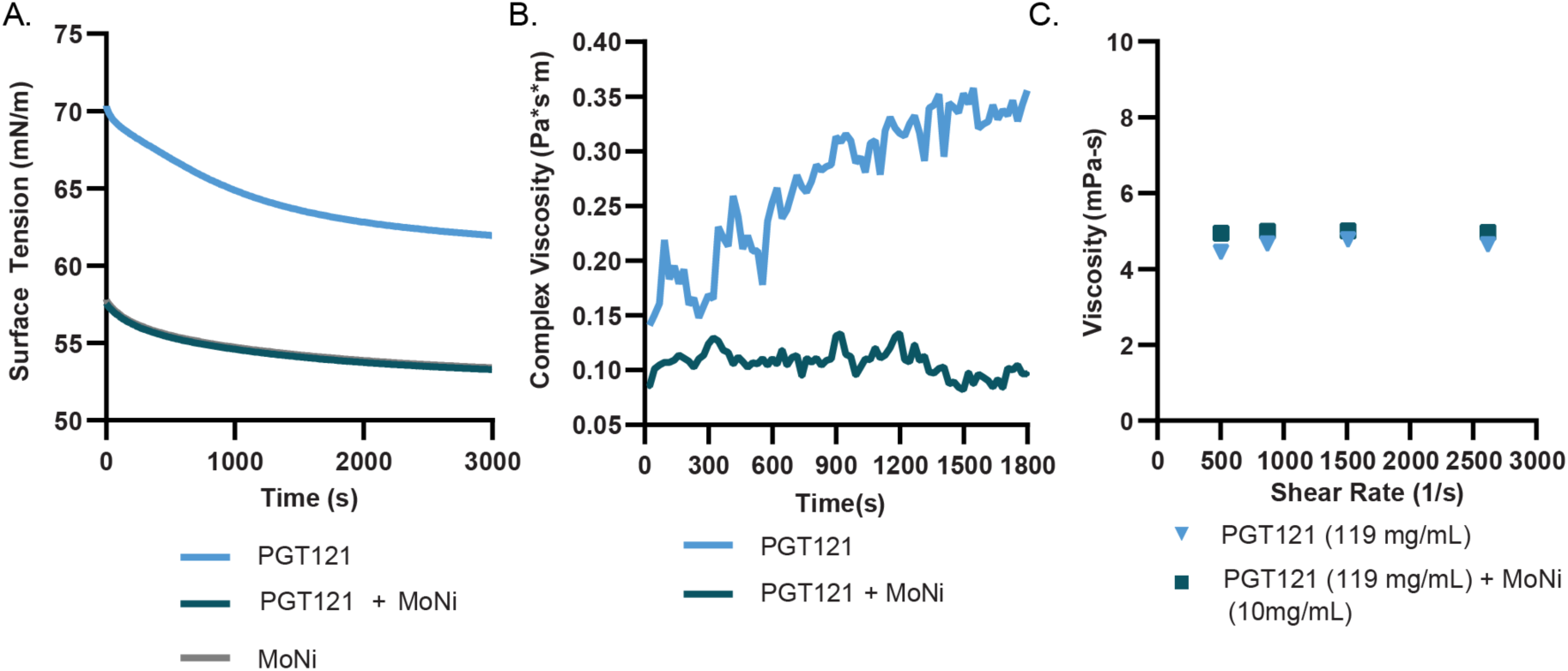
Interfacial interactions of the MoNi polymer excipient in mAb formulations. (A) Comparison of PGT121 (0.2 mg/mL) with and without MoNi (0.1 mg/mL) in time resolved surface tension measurements (n=3). (B) Interfacial rheology measurements (n=3) of PGT121 (0.2 mg/mL) with and without MoNi (0.1 mg/mL). (C) High shear rate viscosity measurements collected at steady state by viscometry experiments for high concentration formulations of PGT121 (119 mg/mL) with and without MoNi (10 mg/mL).

Associative mAb interactions at interfaces leads to the formation of a viscoelastic “skin” and the elasticity of this interface has been shown to correlate strongly with mAb aggregation.^[22] [23]^ Therefore, to further characterize the effects of MoNi on the formulation of PGT121, we probed the interfacial viscoelasticity of these formulations via interfacial shear rheology measurements. These measurements revealed that addition of MoNi to PGT121 formulations reduced the interfacial complex viscosity by 3-fold (from ∼0.3 Pa*s*m to ∼0.1 Pa*s*m; Figure 3B). The high interfacial complex viscosity of the PGT121 formulation without MoNi suggests that protein interactions at the interface are indeed leading to the formation of a gel-like skin at the interface. Consequently, the significant reduction in interfacial complex viscosity upon addition of MoNi indicates that the copolymer excipient reduces interfacial interactions of PGT121, commensurate with the surface tension measurements discussed above that show that the mAbs are precluded from adsorbing to the interfaces. While associative mAb interactions at the interface are reduced upon addition of MoNi, we also sought to determine whether MoNi interacted with the mAbs in the bulk, which could also potentially lead to conformational changes and subsequent aggregation. Using diffusion ordered spectroscopy (DOSY), which is a well-established NMR technique that provides direct evaluation of the diffusion behavior of different species in solution, we demonstrated that the PGT121 mAb and MoNi exhibit distinct diffusion behavior (Figure S1), suggesting that the two species do not interact in the bulk. These results support our hypothesis that MoNi primarily acts to stabilize protein cargo by preventing interfacial aggregation.

With the goal of formulating mAbs at high concentration for SC administration, it was important to verify that formulations with concentrations greater than 100 mg/mL were easily injectable under clinically relevant injection conditions.^[24]^ We measured the viscosity of high-concentration PGT121 (119 ± 7 mg/mL) solutions with and without MoNi (10 mg/mL) at shear rates that are typically achieved during injection through a standard syringe and needle geometry and flow rate. Addition of MoNi at this high concentration did not alter the injection viscosity of the formulation, and both formulations exhibited sufficiently low viscosities to be easily injected (Figure 3C).

### 2.2. MoNi polymer excipient stabilizes mAb formulations in stressed aging assays

The interfacial rheology and surface tension experiments provide strong evidence for preferential adsorption of MoNi to interfaces in formulations with mAbs. To determine whether this behavior also conferred a stability benefit through inhibition of aggregation-nucleating events at the interface, we conducted accelerated aging assays. We compared a stock PGT121 formulation at 55.5 mg/mL (20 mM acetate, 9 wt% sucrose, 0.01 wt% PS80, pH∼5.2) to a formulation concentrated more than 2-fold at 119 ± 7 mg/mL (denoted as high concentration). These formulations with and without MoNi (10 mg/mL) were then packaged in glass vials and subjected to accelerated aging (constant agitation at 50°C; Figure 4A). The monomeric mAb content in these formulations was monitored over four weeks with size exclusion chromatography (SEC; Figure 4B). Within 5 days, both the stock and high concentration PGT121 formulations had lost more than 50% of their monomeric mAb content, indicating significant aggregation of the protein. By day 7, both formulations had macroscopically aggregated, becoming visibly opaque. In contrast, the stock PGT121 formulation comprising MoNi maintained greater than 95% monomeric mAb content through three weeks of continuous stressed aging (Figure 4B). While the high concentration sample comprising MoNi exhibited a small initial decrease in monomer mAb content over the first week of stressed aging, likely arising from our concentration process, this formulation maintained near 70% monomer mAb content over the same three-week period (Figure 4B).

**Figure 4:**
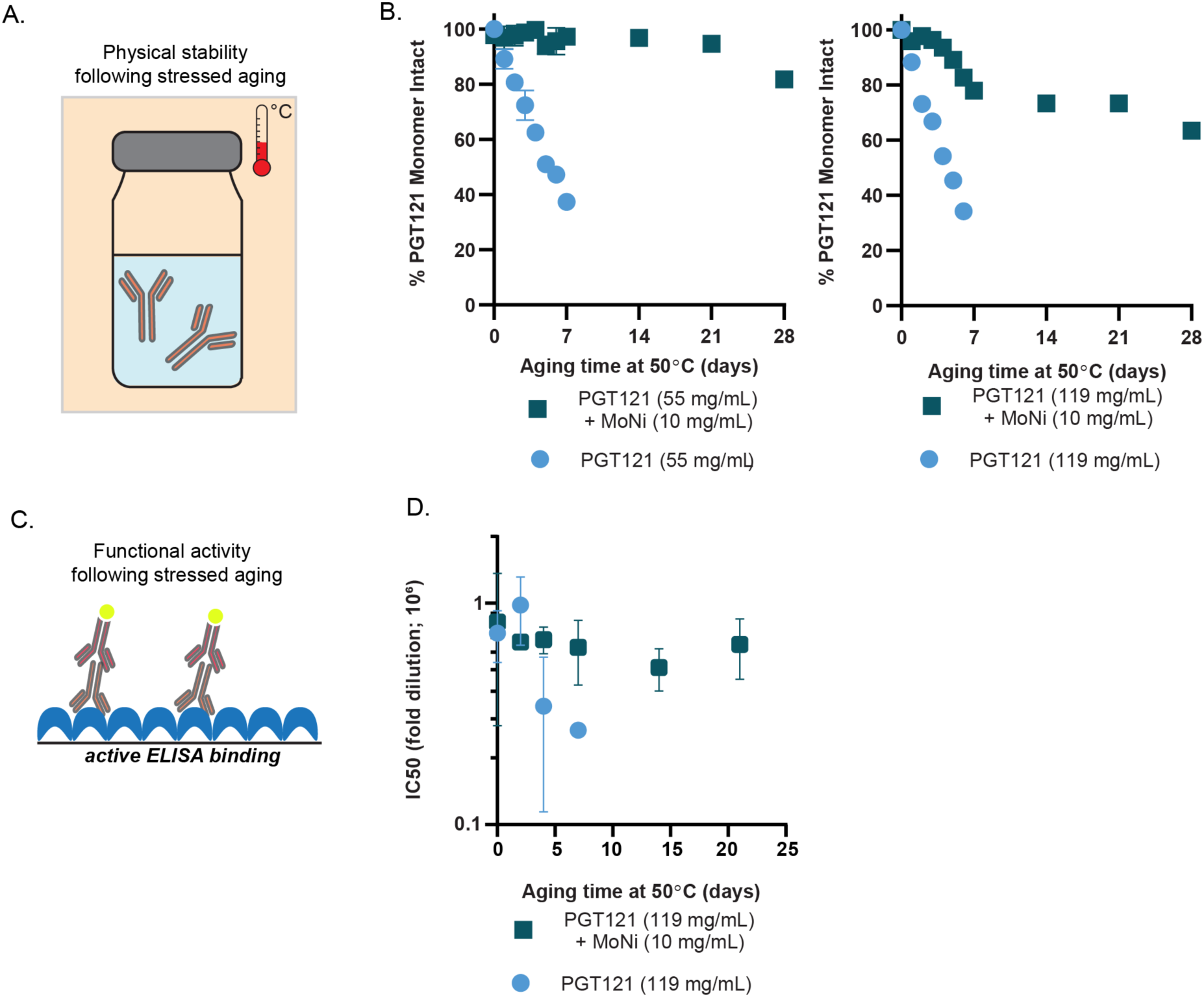
MoNi excipient stabilizes PGT121. (A) Schematic of accelerated aging, whereby formulations are packaged in glass vials, agitated, and heated. (B) Percent monomeric composition of aged samples determined via SEC. (C) Schematic of assaying epitope binding via ELISA. (D) Half maximal inhibitory dilutions of aged formulations with and without MoNi. Samples assayed in duplicate.

For biopharmaceuticals, preservation of monomeric structure is necessary but not sufficient for maintaining therapeutic activity. Critical to the success of antibody-based therapeutics is the conservation of structural domains in the antibody’s binding epitopes. While the MoNi excipient prevented formation of aggregates as determined by SEC, we next sought to determine whether binding epitope activity (as a proxy for therapeutic efficacy) was preserved through continuous stressed aging. We used an enzyme-linked immunosorbent assay (ELISA) to determine whether samples subjected to stressed aging retained binding activity, and thus conformational fidelity, in both their Fab and Fc domains (Figure 4C). Half maximal inhibitory dilutions (IC_50_) were used to compare the functional potency of the high concentration formulations with and without MoNi (10 mg/mL) subjected to stressed aging. In the absence of MoNi, the PGT121 mAbs rapidly lost functionality, losing more than 65% of their potency after just five days of stressed aging (Figure 4E). In contrast, the addition of MoNi significantly enhanced formulation stability and the PGT121 mAbs retained more than 75% of their original potency through 21 days of continuous stressed aging.

Taken together, these data indicate that the addition of the MoNi excipient to high concentration mAb formulations confers a substantial stability benefit by precluding mAb adsorption to the interfaces, thereby preventing aggregation events and maintaining binding activity under accelerated aging conditions.

### 2.3. Pharmacokinetic profile of MoNi-stabilized high concentration PGT121 formulation

To further demonstrate the potential of MoNi as an excipient for mAb formulations, we sought to confirm that it functions as an inactive ingredient by conducting a pharmacokinetic study in rodents. We administered 1.5 mg of PGT121 in transgenic SCID mice with humanized FcRn receptors (n=6/group; B6.Cg-Fcgrt^tm1Dcr^ Prkdc^scid^ Tg[FCGRT]32Dcr/DcrJ; Jackson Labs No. 018441) via either SC injection of 15 μL of MoNi-stabilized (1 wt%) high concentration PGT121 (119 mg/mL) or IP injection of 300 μL of low concentration PGT121 (5 mg/mL). Serum was collected for three weeks post administration to be analyzed for systemic PGT121 concentration via ELISA (Figure 5A,B).

**Figure 5:**
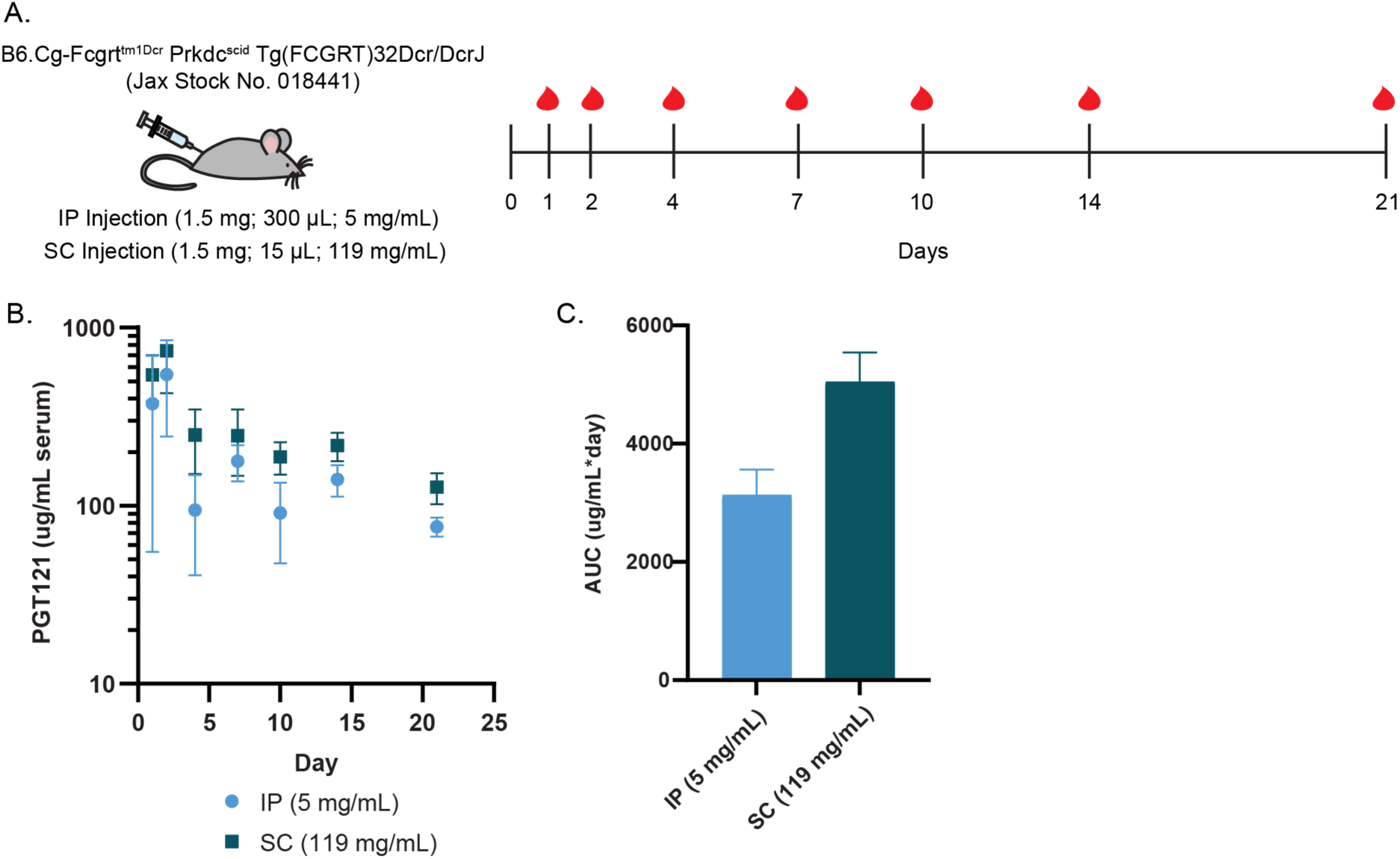
Pharmacokinetics of MoNi-stabilized PGT121. (A) Experimental scheme, indicating injection dosing and blood collection timeline. (B) Serum concentration of PGT121 over time, as determined by ELISA (n=6/group). (C) Area under the curve analysis of ELISA data.

Quantification of serum concentration over time as well as total PGT121 exposure (area-under-the-curve; AUC) demonstrated comparable pharmacokinetics and bioavailability between the two routes of administration and formulations (Figure 5C). We did not observe a therapeutic difference in terms of serum titer nor bioavailability of PGT121 when administered in a MoNi-stabilized, high concentration formulation. Moreover, no acute weight loss (a common metric for treatment related toxicity) was observed, corroborating previous characterization of MoNi’s biocompatibility and tolerability (Figure S2).^[14, 15]^

Overall, the addition of MoNi to high concentration PGT121 mAb formulations significantly improves their stability without altering pharmacokinetics, potentially enabling the development of clinically relevant SC formulations

## 3. Conclusion

In this study, we report on the use of MoNi, an amphiphilic copolymer of acryloylmorpholine and *N*-isopropylacrylamide, as a promising excipient to stabilize high concentration mAb formulations. With an increasing market share and over 64 therapeutic mAbs approved by the FDA,^[25]^ biopharmaceuticals are becoming the predominant class of therapeutics. Unfortunately, producing stable biopharmaceutical formulations at relevant concentrations remains a challenge, as increasing formulation concentration increases the likelihood of aggregation. We had previously developed the copolymer excipient MoNi ^[15]^ for stabilization of insulin, a small protein therapeutic. We hypothesized that MoNi would confer similar stability benefits to mAb formulations by competitively occupying interfaces, such as the air-water interface, reducing the likelihood of interfacially-mediated aggregation of a well-studied anti-HIV bnAb in formulation. Surface tension and interfacial rheology experiments of the PGT121 mAb formulations containing MoNi indicated that the copolymer excipient preferentially adsorbed to the air-water interface, significantly reducing surface tension, and preventing the formation of a viscoelastic gel-like skin of aggregated mAb at the interface. Further, both low (55.5 mg/mL) and high (119 mg/mL) concentration PGT121 solutions formulated with MoNi (10 mg/mL) almost completely maintained monomeric mAb content for over three weeks when subjected to stressed aging by constant agitation at 50°C. In contrast, standard mAb formulations completely aggregated in less than a week under these conditions. Moreover, the ELISA binding activity, which is a good proxy for therapeutic efficacy, was preserved over these same timeframes. Indeed, high concentration PGT121 formulations containing MoNi maintained greater than 75% of their binding activity for more than three weeks of constant stressed aging, whereas standard formulations lost over 65% of their binding activity in only 5 days under the same conditions.

When used as a simple “drop-in” excipient, MoNi does not alter the viscosity of high concentration mAb formulations, as measured by high shear rate viscometry reflective of injection using a standard syringe and needle. Additionally, we demonstrated that the addition of MoNi as an excipient does not alter the pharmacokinetic profile nor the bioavailability of PGT121 in a humanized FcRn mouse model when administered as part of a stabilized high PGT121 concentration formulation injected SC compared to a standard low PGT121 concentration (5 mg/mL) formulation injected IP.

Together, the facile synthesis of MoNi with a scalable controlled radical polymerization technique, its high degree of biocompatibility and low toxicity,^[14, 15]^ and its demonstrated ability to enhance the stability of high concentration biopharmaceutical formulations without altering their pharmacokinetics demonstrate its potential as a simple “drop-in” excipient to improve biopharmaceutical formulations. Enabling stable, high concentration biopharmaceutical formulations can facilitate moving away from IV administration of these drugs in certain applications and more broadly contribute to improved shelf life and a reduction in cold chain dependence.

## 5. Experimental Section

### Materials

Solvents *N,N*-dimethylformamide (DMF; >99.7%; HPLC grade, Alfa Aeser), ethanol (EtOH; >99.5%; Certified ACS, Acros Organics), acetone (>99.9%; Sigma-Aldrich, HPLC Grade), hexanes (>99.9%; Thermo Fisher Scientific, Certified ACS), ether (anhydrous, >99%; Sigma-Aldrich, Certified ACS), and CDCl_3_ (>99.8%; Acros Organics) were used as received. 4-acryloylmorpholine (MORPH, >97%; Sigma-Aldrich) was filtered with basic alumina before use. *N-*isopropylacrylamide (NIP, >99%; Sigma-Aldrich) was used as received. RAFT CTA 2-cyano-2-propyl dodecyl trithiocarbonate (2-CPDT; >97%; Strem Chemicals) was used as received. Initiator 2,2-azobis(2-methyl-propionitrile) (AIBN; >98%; Sigma-Aldrich) was recrystallized from methanol (>99.9%; Thermo Fisher Scientific, HPLC grade) and dried under vacuum before use. Z group removing agents lauroyl peroxide (LPO; 97%; Sigma-Aldrich) and hydrogen peroxide (H_2_O_2_; 30%; Sigma-Aldrich) were used as received. PGT121 monoclonal antibody was provided by Just-Evotec Biologics, Inc. (Seattle, WA, USA) in collaboration with the Bill and Melinda Gates Foundation. PGT121 was supplied at 55.5 mg/ml in 20 mM acetate buffer with 9% (w/v) sucrose, 0.01% PS 80, pH 5.0.^[20]^ 9e9 anti-idiotype monoclonal antibody was synthesized by the Protein Production Facility (PPF) funded by the Bill and Melinda Gates Foundation, using a plasmid provided by Dr. John Mascola at the Vaccine Research Center (VIRC), a division of the National Institute of Allergy and Infectious Diseases (NIAID) in the US National Institutes of Health (NIH).

### SEC characterization

M_n_, M_w_, and Ð for MoNi was determined via SEC implementing PEG standards (American Polymer Standards Corporation) after passing through two SEC columns [inner diameter, 7.8 mm; Mw range, 200 to 600,000 g mol^-1^ ; Resolve Mixed Bed Low divinylbenzene (DVB) (Jordi Labs)] in a mobile phase of DMF with 0.1 M LiBr at 35°C and a flow rate of 1.0 ml min^−1^ [Dionex UltiMate 3000 pump, degasser, and autosampler (Thermo Fisher Scientific)].

### NMR characterization

NMR spectra were recorded in CDCl_3_ on a Varian 600 MHz and δ values are given in parts per million (ppm).

### Synthesis of MoNi

MoNi was synthesized via reversible addition fragmentation transfer (RAFT), as described previously.^[13]^ Briefly, MORPH (645 mg, 4.57 mmol, 41.5 eq.), NIP (105 mg, 0.93 mmol, 8.5 eq.), 2-CPDT (38 mg, 0.11 mmol, 1 eq.), and AIBN (3.6 mg, 0.02 mmol, 0.2 eq.) were combined and diluted with DMF to a total volume of 2.25 ml [33.3 (w/v) vinyl monomer concentration] in an 8-ml scintillation vial equipped with a PTFE septa. The reaction mixture was sparged with nitrogen gas for 10 min and then heated for 12 hours at 65°C. To remove the Z terminus of the resulting polymer, AIBN (360 mg, 2.2 mmol, 20 eq.) and LPO (88 mg, 0.22 mmol, 2 eq.) were added to the reaction mixture, which was then sparged with nitrogen gas for 10 min and heated for 12 hours at 90°C. Z group removal was confirmed by the ratio of the refractive index to ultraviolet (310 nm) intensity in SEC analysis. Resulting polymers were precipitated three times from ether, dried under vacuum overnight, and characterized. SEC: M_n_ = 3059 Da, M_w_ = 3386 Da, and Ð = 1.11.

### Dynamic Light Scattering

DLS measurements of MoNi, prepared in Milli-Q water at 1, 5, 25, 50 and 100 mg/mL, were recorded on a DynaPro Plate Reader II, Wyatt Technology (n=5 acquisitions, average reported).

### Time-resolved surface tension measurements

Time-resolved surface tension of the air–solution interface was measured with a platinum/iridium Wilhelmy plate connected to an electrobalance (KSV Nima, Finland). The Wilhelmy plate was partially immersed in the aqueous solution in a Petri dish, and the surface tension of the interface was recorded for 50 min from the formation of a fresh interface. Equilibrium surface tension values (t = 50 min) were reported as these values more closely describe the environment in a stored vial prior to agitation. Samples were diluted in a 20 mM acetate buffer without sucrose and surfactant, pH = 5.0, from stock samples to the desired assay concentrations. The experiment was repeated in triplicate.

### Interfacial Rheology

Interfacial shear rheology was measured using a Discovery HR-3 rheometer (TA Instruments) with an interfacial geometry comprising a Du Noüy ring made of platinum/iridium wires (CSC Scientific, Fairfax, VA, catalog no. 70542000). Before each experiment, the Du Noüy ring was rinsed with ethanol and water and flame treated to remove organic contaminants. The solution chamber consisted of a double-wall Couette flow cell with an internal Teflon cylinder and an external glass beaker. A time sweep was performed at a strain of 1% (within the linear regime) and a frequency of 0.05 Hz (sufficiently low such that the effects due to instrument inertia will not be significant). Interfacial complex shear viscosity was measured for 30 min. Samples were diluted in a 20 mM acetate buffer without sucrose and surfactant, pH = 5.0, from stock samples to the desired assay concentrations. The experiment was repeated in triplicate.

### Viscometry

A Rheosense m-VROC viscometer with a low viscosity chip was used to measure the viscosity at high shear rates representative of injection. Samples were measured from low to high shear rates using a Hamilton syringe. Each data point was collected at steady state.

### Preparation of high concentration PGT121 formulations

PGT121 monoclonal antibody was concentrated via spin filtration (Corning® Spin-X® UF 5kDa MWCO) at 3030 RPM until desired volume recovery. Concentrated formulations were refrigerated for 12 hours before usage in accelerated aging studies. Concentrations of PGT121 formulations were determined by either volume recovery, ELISA, Nanodrop (Thermo Scientific) or aqueous SEC-UV (PBS buffer, 300 ppm sodium azide), depending on the assay. For aqueous SEC-UV, PGT121 aliquots were diluted 133X in Milli-Q water and concentration was determined by comparing the area under the curve of the traces obtained from SEC using a Dionex UltiMate 3000 VWD at 280 nm (Thermo Scientific). For nanodrop, the protein concentration was compared to a stock standard at 55.5 mg/mL. For ELISA, concentrations were interpolated by fitting a four-parameter dose-response curve (variable slope) in GraphPad Prism 9. The high concentration PGT121 formulation is reported as the mean ± standard deviation of the concentration determined by these methods for several preparations of these formulations.

### In vitro stability assay

150 μL of PGT121 solutions (55.5 mg/mL, 119 mg/mL) were diluted with 7.5 μL of MoNi in the formulation buffer at 21 mg/ml to reach a final excipient concentration of 0.1 wt.%. These formulations were agitated in glass vials at 200 rpm on an orbital shaker plate in an incubator at 50 °C. 5 μL aliquots were removed every 24 hours and stored at 4 °C prior to analysis. Percent monomeric composition and the functional binding activity of aged samples were compared to unaged samples via SEC and ELISA, respectively, as described above.

### PGT121 ELISA

The capture antibody, 9e9, was coated at 2 μg/ml in phosphate buffered saline (PBS) (25 μg per well) on Corning 96-well high binding flat-bottom half-area microplates (Fisher Scientific) and incubated at 4°C overnight. Blocking buffer used was 2% non-fat dry milk (NFDM) in PBS, assay buffer used was 2% bovine serum albumin (BSA) in PBS, and wash buffer was PBS-T (0.05% Tween 20). Plates were washed twice and blocked with 125 μL of blocking buffer and incubated for 1 hr at room temperature. Sample dilutions and PGT121 standards were prepared in assay buffer. After blocking, assay buffer was removed from the plate before adding 50 μL of each sample and incubating for 1 hr at room temperature. Plates were washed five times before adding 50 μL per well of the secondary antibody, Peroxidase AffiniPure F(ab’)_2_ Fragment Goat Anti-Human IgG, Fcγ fragment specific (Jackson Immunoresearch, 109-036-008, RRID: AB\_2337591), diluted 1:5000 in assay buffer. After incubating for 1 hr at room temperature, plates were washed 10 times and then developed by adding 50 μL of TMB ELISA Substrate (High Sensitivity) (Abcam, ab171523). Development was stopped after 2.5 min by adding 1 N hydrochloric acid. PGT121 concentration was measured by absorbance intensity at 450 nm using a Synergy H1 Hybrid Multi-Mode Plate Reader (BioTek). Each plate contained a 15-point standard curve assayed in duplicate which was used to interpolate PGT121 concentrations by fitting a four-parameter dose-response curve (variable slope) in GraphPad Prism 9. Stressed aging samples were diluted 1:10000 and then serially diluted by 4x to create a 7-point curve assayed in duplicate for each sample. IC_50_ values were determined by fitting a four-parameter dose-response curve (variable slope) in GraphPad Prism 9. Mean value and standard deviation reported.

### In vivo pharmacokinetic study

Animal studies were conducted in accordance with the guidelines for care and use of laboratory animals under protocols approved by the Stanford Institutional Animal Care and Use Committee (IACUC). Eight week old female B6.Cg-Fcgrt^tm1Dcr^ Prkdc^scid^ Tg(FCGRT)32Dcr/DcrJ (The Jackson Laboratory, Stock No. 018441) mice were administered PGT121 antibody (1.5 mg/mouse) via IP or SC injection under brief isoflurane anesthesia (n=6/group). This mouse strain was chosen because it is specifically engineered to allow for the study of the human IgG antibody pharmacokinetics.^[26] [27]^ The polymer-stabilized high concentration SC formulation was prepared with 102 mg/ml PGT121 (as determined by ELISA) and 1 wt% MoNi (15 μL injection volume), and the low concentration IP formulation was prepared at 5 mg/ml (300 μL injection volume) in the same buffer in which the antibody was provided. Blood samples were collected at 24 hours, 48 hours, and then days 4, 7, 10, 14, and 21 post-injection for analysis of PGT121 serum concentration via ELISA.

### Statistics

All results are expressed as mean ± standard deviation (SD) unless specified otherwise and analyzed using Graphpad Prism 9.1 (GraphPad Software Inc, USA).

## Ethical Statement

All animal procedures were performed according to Stanford APLAC approved protocols.

## Data Availability

All data supporting the results in this study are available within the article and its Supplementary Information. The broad range of raw datasets acquired and analyzed (or any subsets thereof), which would require contextual metadata for reuse, are available from the corresponding author upon reasonable request.

## Acknowledgements

This work was supported by the Stanford Maternal & Child Health Research Institute through the SPARK Translational Research Program, the Center for Human Systems Immunology with the Bill & Melinda Gates Foundation (OPP1113682; OPP1211043), and the National Institutes of Health (NIAID R01 AI154989). C.M.K. was supported by the Stanford Bio-X William and Lynda Steere Fellowship. J.L.M was supported by the Department of Defense NDSEG Fellowship and by a Stanford Graduate Fellowship. A.I.D. was supported by a Schmidt Science Fellows Award. J.B. was supported by a Marie Curie Postdoctoral Fellowship from the European Union (program H2020, Grant 101030481). A.K.G. is thankful for a National Science Foundation Graduate Research Fellowship and the Gabilan Fellowship of the Stanford Graduate Fellowship in Science and Engineering. Part of this work was performed at the Stanford Nano Shared Facilities (SNSF), supported by the National Science Foundation under award ECCS-1542152.

