## Supplemental Information for "Stable High-Concentration Monoclonal Antibody Formulations Enabled by an Amphiphilic Copolymer Excipient"

### Supplemental Methods

*Diffusion Ordered Spectroscopy (DOSY)*

A 0.2 mg/ml PGT121 sample in $D_2O$ was prepared by diluting the stock material (55.5 mg/ml PGT121 in 20 mM acetate buffer with 9% (w/v) sucrose, 0.01% PS 80, pH 5.0) in $D_2O$ (Acros Organics) and dialyzing against $D_2O$ in 2000 Da MWCO Slide-A-Lyzer dialysis cassettes (Thermo Scientific) for 24 hours prior to use. MoNi was prepared directly in $D_2O$ at 0.1 mg/mL. $^1H$ two-dimensional DOSY spectra were recorded using a Varian Inova 600 MHz NMR instrument. Magnetic field strengths ranged from 2 to 57 G $cm^{-1}$. The DOSY time and gradient pulse were set at 66.5 ms ($\Delta$) and 2 ms ($\delta$), respectively. All NMR data were processed using MestReNova 11.0.4 software.

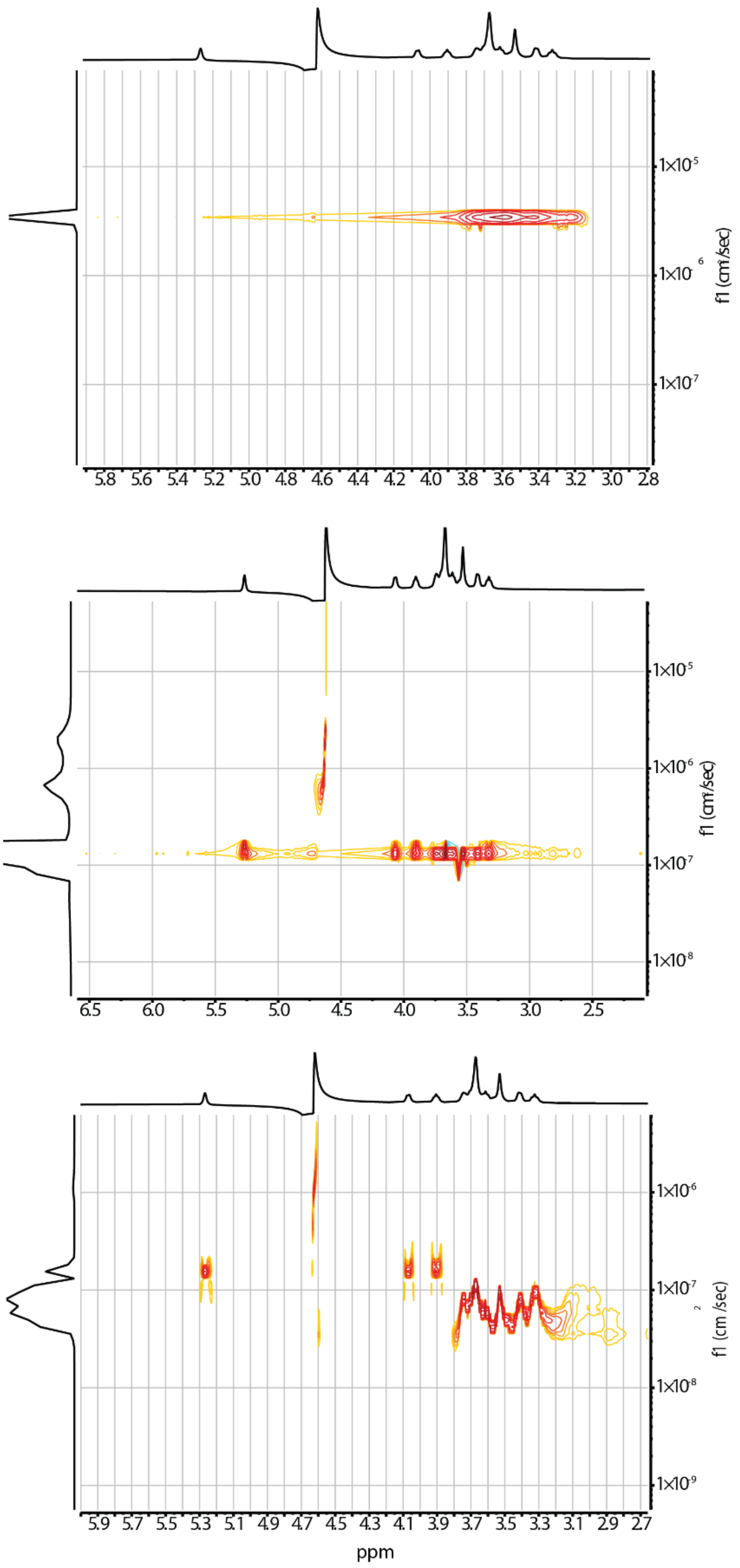


**Figure S1**: Diffusion ordered spectroscopy (DOSY) of (a) MoNi alone (0.1 mg/mL) (b) PGT121 alone (0.2 mg/mL) (c) co-formulation of PGT121 and MoNi at 0.2 mg/mL and 0.1 mg/mL, respectively.

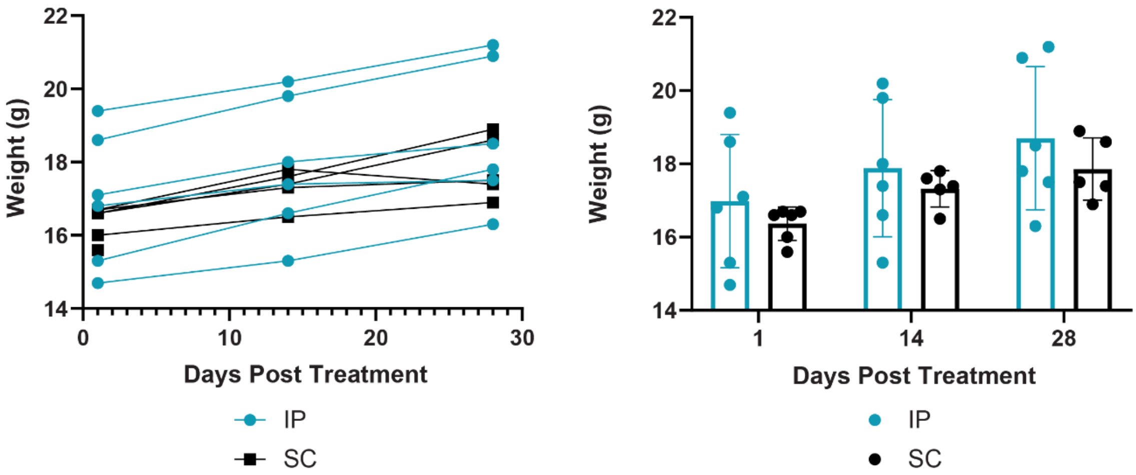


**Figure S2**: Mouse weight over time post treatment
